# *RNA toxicity and perturbation of* rRNA processing in spinocerebellar ataxia type 2

**DOI:** 10.1101/2021.05.07.443200

**Authors:** Pan P. Li, Roumita Moulick, Hongxuan Feng, Xin Sun, Nicolas Arbez, Jing Jin, Leonard O. Marque, Erin Hedglen, H.Y. Edwin Chan, Christopher A. Ross, Stefan M. Pulst, Russell L. Margolis, Sarah Woodson, Dobrila D. Rudnicki

**Affiliations:** Department of Psychiatry and Behavioral Sciences, Division of Neurobiology, Johns Hopkins University School of Medicine, Baltimore, Maryland, USA; T.C. Jenkins Department of Biophysics, Johns Hopkins University, Baltimore, Maryland, USA; Biochemistry Program, School of Life Sciences, The Chinese University of Hong Kong, Hong Kong; Department of Neurology, Johns Hopkins University School of Medicine, Baltimore, Maryland, USA; Department of Neuroscience, Johns Hopkins University School of Medicine, Baltimore, Maryland, USA; Department of Neurology, University of Utah, Salt Lake City, Utah, USA

## Abstract

**BACKGROUND:** Spinocerebellar ataxia type 2 (SCA2) is a neurodegenerative disease caused by expansion of a CAG repeat in *Ataxin-2* (*ATXN2*) gene. The mutant ATXN2 protein with a polyglutamine tract is known to be toxic and contributes to the SCA2 pathogenesis.

**OBJECTIVE:** Here we tested the hypothesis that the mutant *ATXN2* transcript with an expanded CAG repeat (*expATXN2*) is also toxic and contributes to SCA2 pathogenesis.

**METHODS:** The toxic effect of *expATXN2* transcripts on SK-N-MC neuroblastoma cells and primary mouse cortical neurons was evaluated by caspase 3/7 activity and nuclear condensation assay, respectively. RNA immunoprecipitation assay was performed to identify RNA binding proteins (RBPs) that bind to *expATXN2* RNA. Quantitative PCR was used to examine if rRNA processing is disrupted in SCA2 and Huntington disease (HD) human brain tissue.

**RESULTS:** *expATXN2* RNA induces neuronal cell death, and aberrantly interacts with RBPs involved in RNA metabolism. One of the RBPs, transducin β-like protein 3 (TBL3), involved in rRNA processing, binds to both *expATXN2* and expanded *huntingtin* (*expHTT*) RNA *in vitro*. rRNA processing is disrupted in both SCA2 and HD human brain tissue.

**CONCLUSION:** These findings provide the first evidence of a contributory role of *expATXN2* transcripts in SCA2 pathogenesis, and further support the role *expHTT* transcripts in HD pathogenesis. The disruption of rRNA processing, mediated by aberrant interaction of RBPs with *expATXN2* and *expHTT* transcripts, suggest a point of convergence in the pathogeneses of repeat expansion diseases with potential therapeutic implications.

## Introduction

Spinocerebellar ataxia type 2 (SCA2) is an autosomal dominant disorder caused by a CAG repeat expansion in the first exon of the *ATXN2* gene located on chromosome 12q24 ^1^. The repeat is in-frame to encode polyglutamine (polyQ). The signs and symptoms of SCA2 include progressive deterioration in balance and coordination, neuropathies, nystagmus and slow saccadic eye movements, slurred speech and cognitive impairment ^2-5^. SCA2 is the second most common form of autosomal dominant ataxia, with a prevalence of 1-2 cases/10^5^ inhabitants, varying somewhat by ethnicity and geographic location ^2, 3, 6-8^. The highest prevalence of the SCA2 mutation occurs in Cuba (6.57 cases/10^**5**^ inhabitants) ^9^, and is likely a consequence of a founder effect ^10^. SCA2 neuropathology is characterized by a significant loss of cerebellar Purkinje neurons, a less prominent loss of cerebellar granule cells ^11^; marked neuronal loss in the inferior olive, pontocerebellar nuclei, and substantia nigra; degeneration of the thalamus and pons, and thinning of the cerebellar cortex without changes in neuronal density ^11-14^. The normal *ATXN2* allele contains 15 to 32 CAG triplets, while the disease allele typically has 33 to 64 triplets ^15^. The most common disease allele has 37 triplets, and neonatal onset SCA2 cases with over 200 CAG repeats have been reported ^16^. Similar to other CAG repeat diseases, the repeat length in SCA2 is inversely correlated to age of onset ^17, 18^.

Recently, intermediate CAG expansion in *ATXN2* has been associated with a higher risk for amyotrophic lateral sclerosis (ALS) ^19^. Current evidence indicates that neurotoxicity of ATXN2 protein, which is involved in multiple cellular pathways, including mRNA maturation, translation, and endocytosis, is central to SCA2 pathogenesis ^20, 21^. This is supported by data from several SCA2 cell and mouse models expressing mutant ATXN2 protein ^22-24^.

However, multiple laboratories, including ours, have demonstrated an important neurotoxic role for mutant RNA transcripts in CAG/CTG repeat expansion diseases, including myotonic dystrophy type 1 (DM1)^25, 26^, Huntington disease (HD) ^27, 28^, Huntington disease-like 2 (HDL2) ^29^, SCA3 ^30-32^, and SCA8 ^33^. RNA-triggered pathogenic processes are thought to be, at least in part, mediated by aberrant interaction between expanded repeat-containing RNA transcripts and RNA-binding proteins (RBPs) ^34-36^. The basic hypothesis is that expanded CAG/CUG repeats in transcripts form hairpin structures which sequester multiple RBPs and hence prevent the RBPs from performing their normal function in cells ^35^. To add to the pathomechanistic complexity of CAG/CUG repeat diseases, antisense transcripts that span the CAG/CUG repeat regions are also expressed at the DM1 (CAG direction)^37^, HDL2 (CAG direction)^38, 39^, SCA7 (CUG direction)^40^, SCA8 (CUG direction) ^33^ and HD (CUG direction)^41^ loci. We have recently described a transcript expressed antisense to *ATXN2* at the SCA2 locus ^42^, and provided evidence that this antisense *ATXN2* (*ATXN2-AS*) transcript contributes to SCA2 pathogenesis, and potentially to ALS associated with an intermediate repeat expansion at the *ATXN2* locus ^19^. We hypothesized that, in addition to mutant ATXN2 protein and mutant *ATXN2-AS* transcript ^42^, mutant sense *ATXN2* RNA also contributes to SCA2 pathogenesis.

As predicted by this hypothesis, the data presented here demonstrate that sense *expATXN2* transcripts are neurotoxic in cell models in the absence of expression of mutant ATXN2 protein, aberrantly interact with RBPs that are involved in rRNA processing, and lead to disruption of rRNA processing. We demonstrate a similar disruption of rRNA processing in HD patient brain tissue. Similar to findings in other repeat expansion diseases, SCA2 is therefore the fifth neurodegenerative CAG/CTG repeat expansion disease in which pathogenesis is likely a consequence of a combination of expression of mutant protein and bi-directionally expressed mutant RNA.

## Materials and Methods

Description of materials and methods is provided in the supplementary data.

## Results

### The non-translatable expATXN2 transcript is neurotoxic

To confirm the toxicity of *expATXN2* transcripts, we cloned FL *ATXN2* cDNA with 22, 58, or 104 CAG triplets into the 3’ untranslated (UTR) region of *Renilla luciferase* (*Rluc*) cDNA, thereby allowing expression of *expATXN2* RNA transcripts, but preventing ATG-initiated translation of the RNA into FL ATXN2 protein (Fig. 1A). No evidence of expression of an expanded polyglutamine tract was detected by Western blotting using the expanded polyglutamine-specific antibody 1C2 ^43^, confirming that the 3’ UTR cloning approach indeed eliminated detectable ATG-initiated translation of the FL *ATXN2* (Fig. 1B). Caspase 3/7 activity assay showed that overexpression of *Rluc-ATXN2-(CAG)58* or *Rluc-ATXN2-(CAG)104* was significantly more toxic than *Rluc-ATXN2-(CAG)22* in SK-N-MC cells (Fig. 1C). Comparable expression levels of overexpressed transcripts in SK-N-MC cells were confirmed by qPCR (Fig. 1D). However, hairpin-forming expanded CAG repeats can also be translated in the absence of ATG start codon through the mechanism of repeat-associated non-ATG translation (RAN translation) ^44^.

**Fig. 1.**
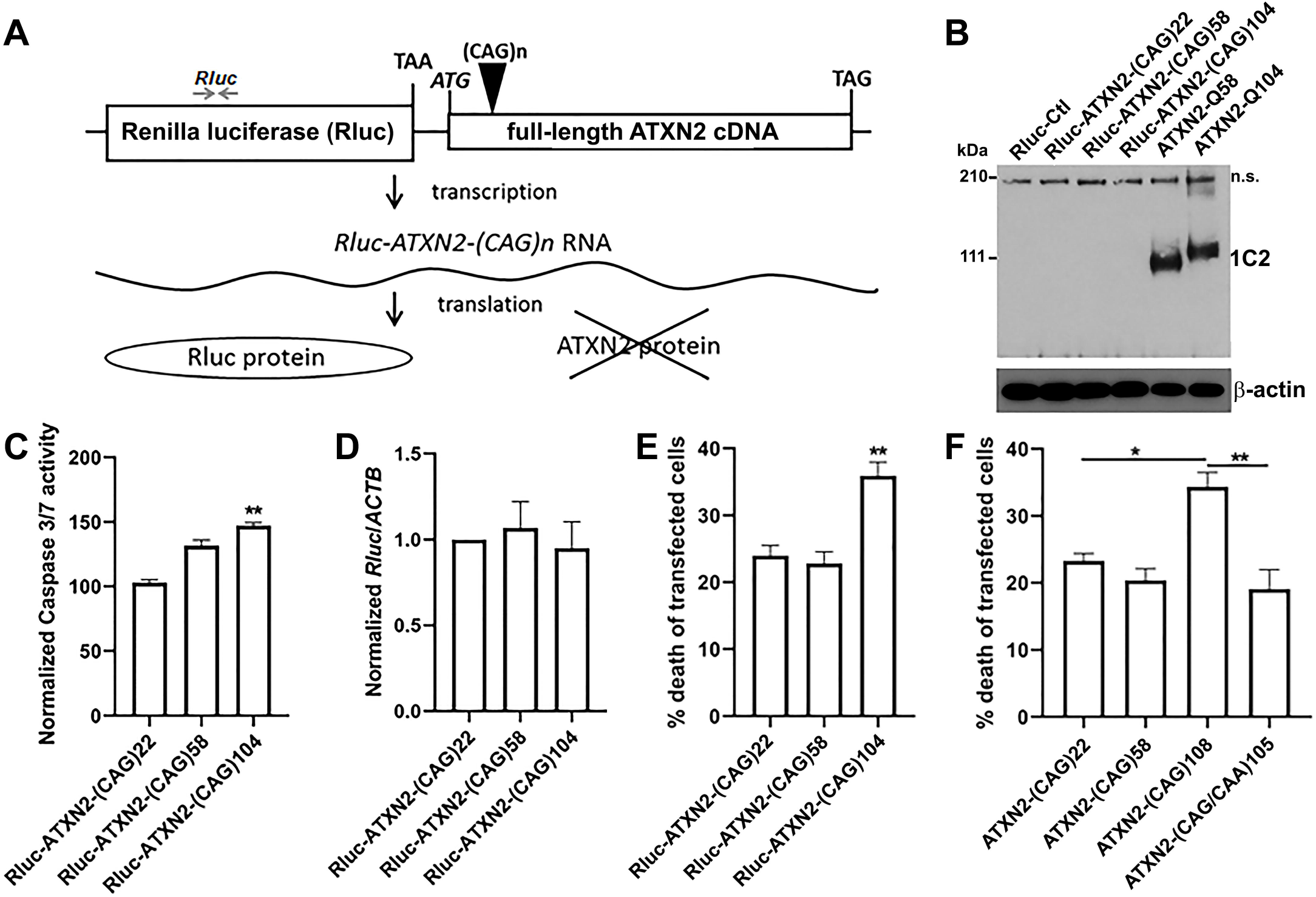
Non-translatable full length *expATXN2* transcript is neurotoxic to SK-N-MC cells. (A) Schematic presentation of the non-translatable full length (FL) *ATXN2* cell model. FL *ATXN2* cDNA was cloned into 3’ UTR region of Renilla luciferase (Rluc) cDNA to prevent its translation. (B) *Rluc-ATXN2-(CAG)n* constructs does not express canonically or RAN translated polyglutamine (polyQ), as confirmed by immunoblotting with polyQ-specific 1C2 antibody. β-actin was used as a loading control. A representative blot was shown. (C) At 72 hours after overexpression, both *Rlu-ATXN2-(CAG)58* and *Rluc-ATXN2-(CAG)104* transcripts are toxic to neuronal-like SK-N-MC cells, as determined by Caspase 3/7 activity assay. The Caspase 3/7 activity in *Rluc-ATXN2-(CAG)22* transfected SK-N-MC cells was normalized to 100. Data were expressed as mean ± SEM from 4 independent samples per condition (N=4); **p<0.01 by Kruskal–Wallis test and Dunn’s multiple comparison test. (D) Comparable expression levels of exogenous *Rluc-ATXN2-(CAG)n* transcripts were confirmed by qPCR. *ACTB* transcript was used as an internal control. Locations of qPCR primers for Rluc were indicated in A. The *Rluc*/*ACTB* ratio in *Rluc-ATXN2-(CAG)22* transfected SK-N-MC cells was normalized to 1. Data were expressed as mean ± SEM from 4 independent samples per condition (N=4); Kruskal– Wallis test. (E) At 48 h after overexpression, *Rluc-ATXN2-(CAG)104* is toxic to primary mouse cortical neurons, as determined by a nuclear condensation assay. (F) *expATXN2* transcript toxicity depends on the repeat’s ability to form toxic hairpin structures. *ATXN2-(CAG)104*, but not the interrupted *ATXN2-(CAG/CAA)105*, is toxic to primary mouse cortical neurons, as indicated by a nuclear condensation assay. Data are expressed as mean ± SEM from 4-8 independent samples per condition (N=4-8). In each sample, 4,000 neurons per condition were analyzed. *p<0.05, **p<0.01 by Kruskal–Wallis test and Dunn’s multiple comparison test.

To exclude the possibility that RAN translation of protein fragments with expanded amino acid tracts leads to neurotoxicity in our SK-N-MC model system, we cloned an *ATXN2* fragment containing a CAG repeat expansion, multiple upstream stop codons, and 150 bp of ATXN2 sequence flanking the repeat (thereby excluding all ATGs) into a vector with tags for each of the three open reading frames (Fig. S1A). There were no detectable protein fragments from any of the three reading frames (Fig. S1B-E), indicating that SK-N-MC cells do not support the RAN translation of *expATXN2* transcripts, and confirming that expression of FL *expATXN2* transcript is sufficient to trigger neurotoxicity even when the transcripts are not translated into proteins.

Consistent with these observations in neuroblastoma cells, overexpression of *Rluc-ATXN2-(CAG)104* triggers neurotoxicity in primary mouse cortical neurons, as measured by nuclear condensation assay (Fig. 1E). *Rluc-ATXN2-(CAG)58* was not toxic in this assay, perhaps reflective of the short time frame of the experiment (nuclear condensation is a later stage event, whereas caspase 3/7 activation occurs at an early stage in cell death), differences in the levels of transcript expression in primary neurons compared to neuroblastoma cell lines, or different sensitivity to transcript-induced toxicity in primary neurons and SK-N-MC cells.

It has been suggested that the CAG repeats form stable hairpin structures ^35^, while CAA interruptions either break hairpin regularity or induce the formation of branched structures ^45^. To examine whether preventing the formation of hairpin structures in *expATXN2* ameliorate its neurotoxicity, we replaced the pure CAG repeat region in the *ATXN2-(CAG)104* with a fragment of heavily interrupted CAG/CAA triplets, to obtain the *ATXN2-(CAG/CAA)105* construct. Inserting interruptions abolished *expATXN2* toxicity in primary mouse cortical neurons (Fig. 1F), suggesting that the secondary hairpin structure adopted by the pure CAG repeat may be critical for neurotoxicity.

### Full-length expATXN2 transcripts form RNA foci in SCA2 cell and mouse models, and in one human SCA2 brain

Repeat-containing mutant transcripts form RNA foci in all CUG/CAG diseases in which RNA neurotoxicity has been demonstrated to contribute to pathogenesis ^46-48^. We therefore sought to detect similar foci in SCA2 models and human brain by fluorescence *in situ* hybridization (FISH). CAG RNA foci were absent in SK-N-MC neuroblastoma cells that overexpress *GFP* alone (Fig. 2A) or a FL ATXN2 construct modified to have only one CAG triplet (*GFP-ATXN2Q1*, Fig. 2B), and were only rarely detected in cells overexpressing FL normal *ATXN2* (*nATXN2*) transcripts with 22 triplets (*GFP-ATXN2Q22*, Fig. 2C). Foci were much more abundant in cells overexpressing full-length (FL) expanded *ATXN2* (*expATXN2*) transcripts with 58 or 104 CAG triplets (*GFP-ATXN2Q58* or *GFP-ATXN2Q104*, Fig. 2D and 2F), as quantified in Fig. 2E. The foci are resistant to DNase treatment and are degraded by RNase treatment (Fig. 2G and 2H).

**Fig. 2.**
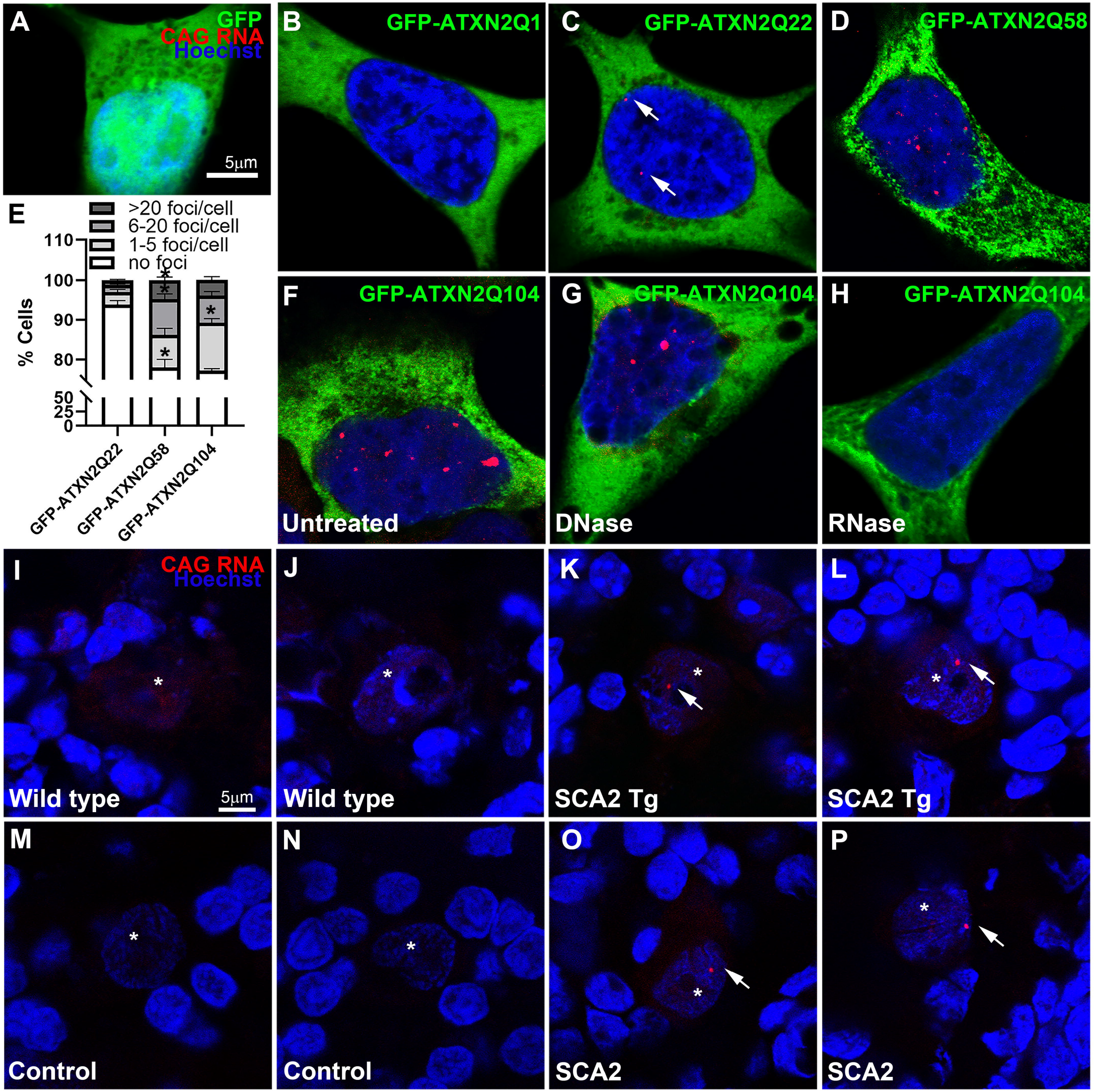
*expATXN2* transcripts form RNA foci. (A-H) Exogenous *expATXN2-AS* transcripts form nuclear CAG RNA foci in SK-N-MC neuroblastoma cells. *GFP-ATXN2-(CAG)58 or 104* (D and F) transcripts formed frequent foci, while RNA foci were occasionally detected in *GFP-ATXN2-(CAG)22* expressing cells (C) and were absent in cells expressing GFP (A) or *GFP-ATXN2-(CAG)1* (B). (E) The percentage of cells with foci. Data were expressed as mean ± SEM from 3 independent samples per condition (N=3); *p<0.05 by Kruskal–Wallis test and Dunn’s multiple comparison test, compared to *GFP-ATXN2-(CAG)22*. (G-H) *GFP-ATXN2-(CAG)104* RNA foci were resistant to DNase treatment (G) and degraded by RNase treatment (H). (I-L) *expATXN2* transcript forms RNA foci in the cerebellar Purkinje cells of SCA2 transgenic (Tg) mice in which the expression of FL *ATXN2-Q127* cDNA is driven by a Purkinje cell specific Pcp2 promoter ^24^ (K-L). RNA foci were not detected in wildtype control mice (I-J). (M-P) *expATXN2* transcript forms RNA foci in cerebellar Purkinje cells of a human SCA2 brain (O-P), but not in human control cerebella (M-N). Scale bar: 5μm. Arrows point to RNA foci and asterisks indicate the Purkinje cells.

This set of experiments demonstrates that *expATXN2* transcripts form RNA foci, and that the extent of foci formation may at least partially correlate with repeat length. Furthermore, while not detected in wildtype (WT, Fig. 2I-J) mice, *ATXN2* RNA foci are present in cerebellar Purkinje neurons of SCA2 transgenic mice (Fig. 2K-L) which express FL *ATXN2* with 127 CAG triplets specifically in Purkinje neurons ^22^. Finally, out of the five human postmortem brains available for this study, *ATXN2* RNA foci were detected in cerebellar Purkinje cells in one brain (H1 case, Table 1) that had 38 triplets for the mutant allele (Fig. 2O-P), but not in the control human brains (Fig. 2M-N). RNA foci may be only a hallmark for RNA toxicity, and whether RNA foci are toxic or not remains to be further determined.

**Table 1.**
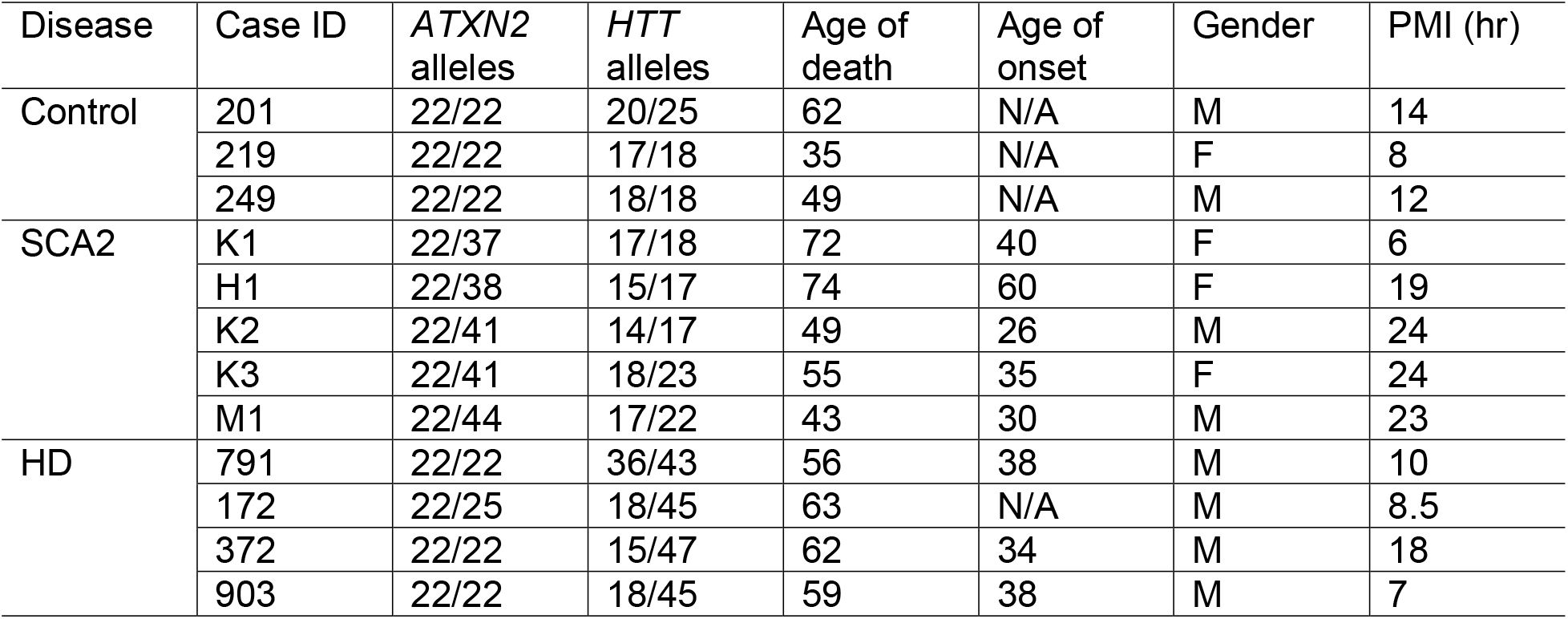
Human control and patient brain information.

| Disease | Case ID | *ATXN2* alleles | *HTT* alleles | Age of death | Age of onset | Gender | PMI (hr) |
| --- | --- | --- | --- | --- | --- | --- | --- |
| Control | 201 | 22/22 | 20/25 | 62 | N/A | M | 14 |
|  | 219 | 22/22 | 17/18 | 35 | N/A | F | 8 |
|  | 249 | 22/22 | 18/18 | 49 | N/A | M | 12 |
| SCA2 | K1 | 22/37 | 17/18 | 72 | 40 | F | 6 |
|  | H1 | 22/38 | 15/17 | 74 | 60 | F | 19 |
|  | K2 | 22/41 | 14/17 | 49 | 26 | M | 24 |
|  | K3 | 22/41 | 18/23 | 55 | 35 | F | 24 |
|  | M1 | 22/44 | 17/22 | 43 | 30 | M | 23 |
| HD | 791 | 22/22 | 36/43 | 56 | 38 | M | 10 |
|  | 172 | 22/25 | 18/45 | 63 | N/A | M | 8.5 |
|  | 372 | 22/22 | 15/47 | 62 | 34 | M | 18 |
|  | 903 | 22/22 | 18/45 | 59 | 38 | M | 7 |

### expATXN2 transcripts aberrantly interact with RNA binding proteins (RBPs)

We next examined whether the neurotoxicity of *expATXN2* transcript is mediated by aberrant *expATXN2* RNA-RBP interactions. We performed an *in vitro* biotinylated *ATXN2* RNA pull-down assay (Fig. 3A) and identified by mass spectrometry (MS) a total of 57 RBPs that preferentially bind to the *expATXN2*, compared to the *nATXN2* transcript. Go analysis of functional annotation^49^ and STRING analysis^50^ of the *expATXN2* RBPs are shown in Fig. S2. A selective list of *expATXN2 RBPs* is shown in Table S1. Out of the 57 *expATXN2* RBPs, 40 are localized in the nucleus, with 20 of them in the nucleolus, suggesting that aberrant *expATXN2*-RBP interactions may predominantly occur in the nucleus. Interestingly, among the 20 nucleolar RBPs, 7 of them contain WD40 repeat domains, of which, five (PWP1, TBL3, WDR3, WDR36 and UTP18; Table S1 and Fig. S2B) are components of the small subunit (SSU) processome for ribosomal RNA (rRNA) processing. We therefore became interested in the SSU processome components that were identified as *expATXN2* RBPs. Out of the five SSU components ^51, 52^ that are potential *expATXN2* interactors, we selected TBL3 (transducin β-like protein 3) for further analysis, as we were interested in RNA mediated disease mechanisms shared by both SCA2 and HD, and by the same method, TBL3 appeared to interact with the expanded *Huntingtin* (*expHTT*) transcript as well (Fig. 3B-C), and has a relatively greater number of peptide hits and percentage of protein coverage, compared with other SSU components identified by MS (Table S1), though the number of peptide hits does not always imply stronger interaction ^53^.

**Fig. 3.**
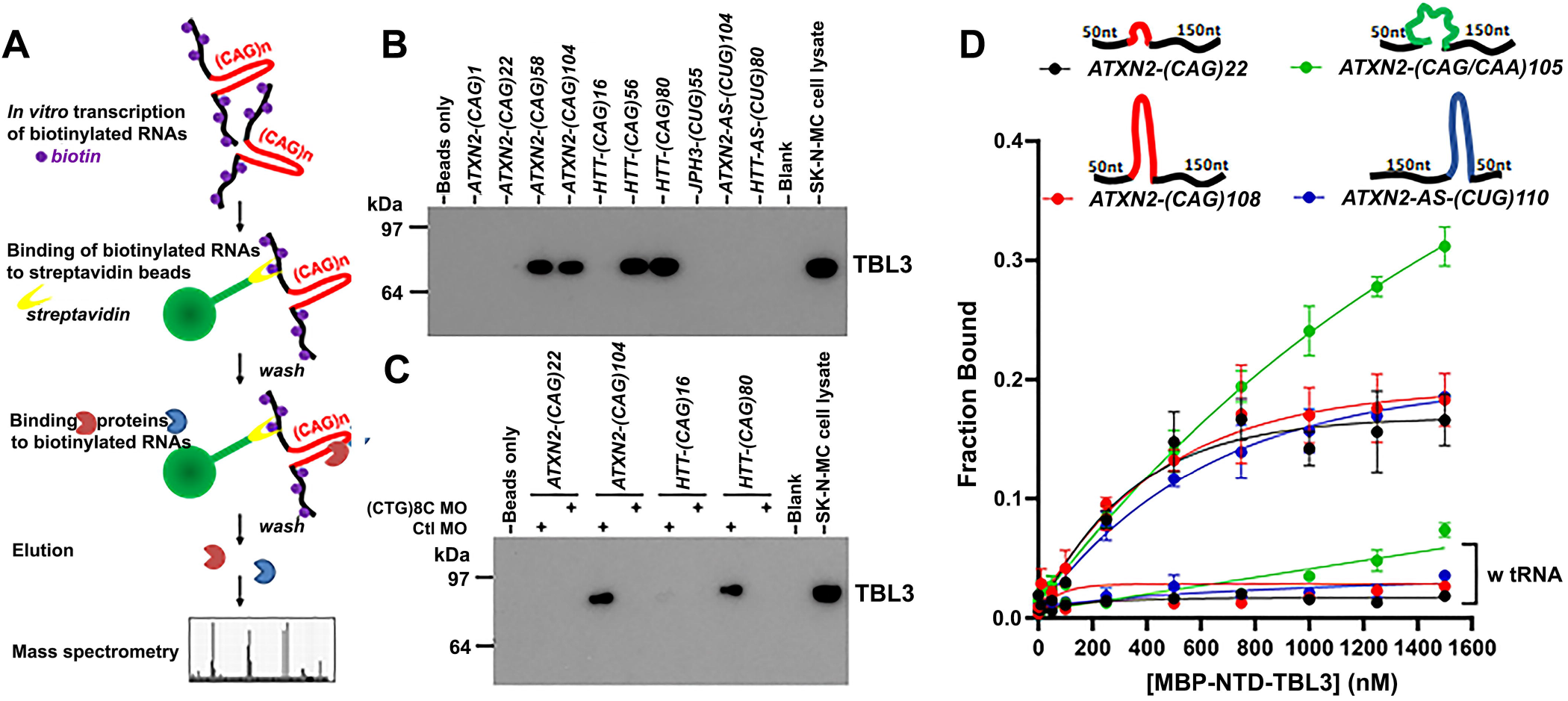
TBL3 aberrantly interacts with *expATXN2* and *expHTT* transcripts. (A) Schematic illustration of the biotinylated RNA pull-down procedure. (C) TBL3 interacts with biotinylated *expATXN2* (with 58 or 104 CAG repeats) and *expHTT* (with 56 or 80 CAG repeats) RNAs in an *in vitro* biotinylated RNA pull-down assay. The interaction of TBL3 is specific for expanded CAG repeats, as no binding between TBL3 and CUG repeats in *JPH3-(CUG)55, ATXN2-AS-(CUG)104* or *HTT-AS-(CUG)80* was observed. (D) Interaction between TBL3 and *expATXN2* or *expHTT* is dependent on the CAG repeat region. Incubation with (CTG)8C, but not control Morpholino (MO), abolished the binding of TBL3 to *ATXN2-(CAG)104* or *HTT-(CAG)80* transcript. SK-N-MC cell lysate was used as a positive control. N=3 independent experiments and representative blots were shown. (E) Nitrocellulose filter-binding analysis of MBP-NTD-TBL3 binding to *ATXN2-(CAG)22, 108, ATXN2-(CAG/CAA)105*, and *ATXN2-AS-(CUG)110* RNAs. The circles represent the mean fraction RNA bound to MBP-NTD-TBL3 in the absence or presence of competitor tRNA (w tRNA), respectively. Schematic presentation of the transcripts with various repeats and lengths of flanking regions are shown. Mean and SEM are shown; n ≥ 3 independent trials. The fits through the data are from non-linear regression analysis of the binding curves to a Scatchard plot. In the absence of competitor tRNA, the K_D_ obtained for *ATXN2-(CAG)22, ATXN2-(CAG)108, ATXN2-(CAG/CAA)105*, and *ATXN2-AS-(CUG)110* were 350 nM, 420 nM, 2.1μM and 650 nM respectively.

### *TBL3 binds to expanded CAG repeats* in vitro

We performed additional RNA pull down experiments and western blots to confirm that TBL3 interacts with *expATXN2 in vitro* (Fig. 3B). To test whether the interaction is disease-specific, we also included *expHTT* transcripts, associated with HD, the most prevalent and most studied CAG repeat disease ^27, 28, 41, 54^. Studies from multiple laboratories, including ours, support the idea that RNA neurotoxicity contributes to HD ^27-29, 47^ We confirmed that TBL3 interacts *in vitro* with expanded CAG repeats flanked with either *ATXN2* or *HTT*-specific sequence (Fig. 3B), but not with expanded CUG repeats flanked with either antisense *ATXN2* (*ATXN2-AS*; ^42^), antisense *HTT* (*HTT-AS*; expressed on HD locus)^41^, or junctophilin-3 (*JPH3)* flanking sequence ^29^ (Fig. 3B). To further confirm that the interaction between TBL3 and *expATXN2* and *expHTT* was dependent on the CAG repeat, we pre-incubated *expATXN2* and *expHTT* transcripts with (CTG)8C Morpholino (MO), which we have previously established hybridizes to CAG repeat expansions ^55^. The pretreatment with (CTG)8C prevented TBL3 from binding to either transcript *in vitro* and provided further evidence that both *expATXN2*-TBL3 and *expHTT*-TBL3 interactions are dependent on the presence of an expanded CAG repeat (Fig. 3C). Taken together, these data indicate that TBL3 binds to expanded CAG repeats independent of flanking sequence.

To investigate whether TBL3 binds to expanded CAG repeats independently of other cellular proteins, we purified the TBL3 N-terminal RNA binding domain as a fusion with maltose binding protein (MBP-NTD-TBL3) and measured its binding with *expATXN2* transcripts using an *in vitro* nitrocellulose filter binding assay ^56^. The isolated TBL3 NTD associated with *ATXN2* CAG RNA, with K_D_ =350 nM and 420 nM for *ATXN2-(CAG)22* and *ATXN2-(CAG)108*, respectively (Fig. 3D). Although overall binding was weak, the yeast homolog of TBL3, Utp13, binds pre-rRNA as a tetramer with other UtpB complex proteins. Therefore, the weak affinity of the isolated MBP-NTD-TLB3 for *expATXN2* may be due to the absence of its normal binding partners. The *in vitro* binding reactions saturated approximately 20-30% of refolded *ATXN2* RNA, suggesting that a fraction of the *ATXN2* RNA is unable to refold into a conformation that is competent to bind TBL3.

Despite its weak affinity for RNA, MBP-NTD-TBL3 bound *ATXN2* CAG repeats more strongly than control RNAs, including the *ATXN2-AS-(CUG)110* transcript (K_D_=650 nM), and a CAG repeat containing CAA interruptions, *ATXN2-(CAG/CAA)105* (K_D_=2.1 μM). This preference for continuous CAG repeats raised the possibility that TBL3 recognizes the hairpin structure of CAG repeat RNA. To test this idea, the filter binding assays were also carried out in the presence of a competitor yeast tRNA, which is expected to be structured under our assay conditions. The tRNA competitor abolished the interaction between MBP-NTD-TBL3 and *ATXN2* or *ATXN2-AS* transcripts (Fig. 3D), consistent with the idea that TBL3 binding depends on the structures of *ATXN2* CAG repeats.

### The effect of TBL3 reduction on 45S pre-rRNA level and processing

Depletion of UTP13, the yeast homolog of TBL3, increases the steady-state level of unprocessed 35S pre-rRNA in yeast ^57^. We therefore hypothesized that, although the interaction between *expATXN2* and TBL3 may not be direct and likely involves other proteins, sequestration of TBL3 in a complex that interacts with *expATXN2* may disrupt its normal function and affect rRNA maturation. We therefore predicted that knockdown of TBL3 in cells, mimicking its sequestration, would increase the level of unprocessed 45S pre-rRNA, the human counterpart of yeast 35S pre-rRNA. Three individual siRNAs were used to knock down TBL3 in HEK293T cells in order to minimize the possibility of alternative mechanisms of TBL3 reductions through off-target effects. Each siRNA reduced TBL3 protein level by 50-80% at 72 hours post transfection (Fig. 4A-B). Next, we examined 45S pre-rRNA levels by qPCR using primers against the 5’ external transcribed spacer ^58^ as indicated in Fig. 4C. Knockdown of TBL3 in HEK293T cells using each siRNA increased steady-state 45S pre-rRNA levels (normalized to *ACTB*, Fig. 4D), consistent with a previous study using stable shRNA transfection ^59^. As previously reviewed ^60, 61^, a complex sequence of cleavage steps is required to release the mature RNAs (18S, 5.8S and 28S) from the precursor 45S pre-rRNA. qPCR using primers against 18S rRNA would detect the mature 18S rRNA, unprocessed 45S pre-rRNA, as well as any intermediate rRNAs containing 18S sequence. Similarly, qPCR using primers against 28S rRNA would detect the mature 28S rRNA, the 45S pre-rRNA precursor, as well as intermediate rRNAs that contain the 28S sequence (Fig. 4C). We therefore used the ratios of 18S rRNA/45S pre-rRNA and 28S rRNA/45S rRNA measured by qPCR, as readouts for 18S rRNA maturation and 28S rRNA maturation, respectively. Depletion of UTP13 in yeast has been previously shown to decrease 18S rRNA maturation ^62, 63^. Consistently, we found that knock down of TBL3 in HEK293T cells decreased the ratio of 18S rRNA to 45S pre-rRNA (Fig. 4E), indicating that TBL3 may play a role in 18S rRNA maturation. In addition, 28S rRNA maturation was also decreased after TBL3 knock down (Fig. 4F). We attempted to determine if overexpression of TBL3 has the opposite effect, however, forced expression of TBL3 by itself triggered toxicity and mis-localized the protein into nuclear aggregates (data not shown). Similarly, MBNL1, an RBP previously shown to interact with expanded CAG/CUG (*expCAG/CUG*) transcript also appeared to be toxic when overexpressed or knocked down ^28^, suggesting that expression of certain RBPs must be tightly controlled to maintain their normal function.

**Fig. 4.**
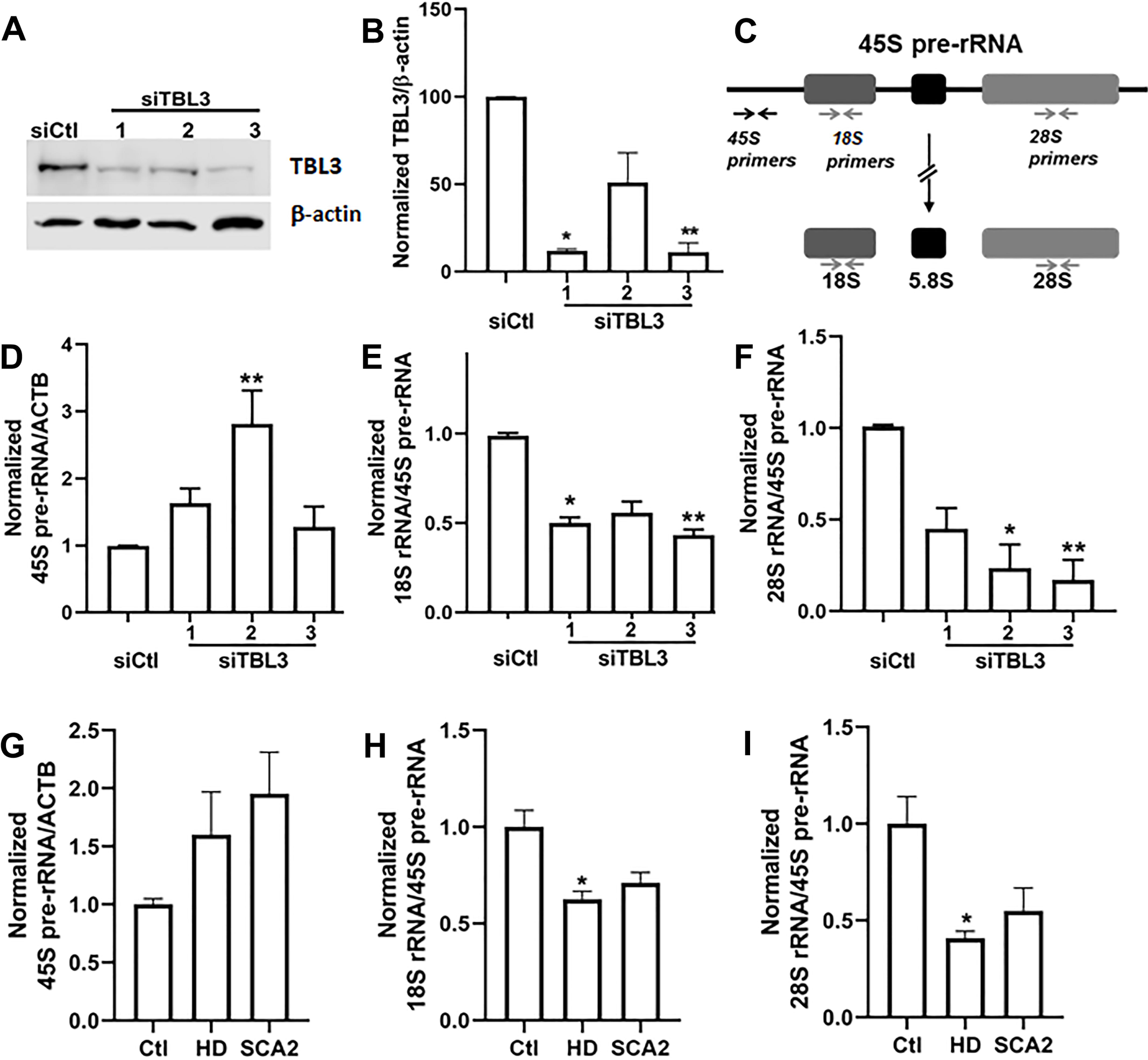
rRNA processing is affected in TBL3 knock-down cells, as well as in SCA2 and HD brains. (A-B) siRNAs against TBL3 (siTBL3), but not control siRNAs (siCtl), efficiently knocked down TBL3 protein expression by 50-80% in HEK293T cells after 72 hours. Representative blots were shown. TBL3/β-actin protein expression in siCtl treated cells was normalized to 100. Data were expressed as mean ± SEM from 3 independent samples per condition (N=3). *p < 0.05 and **p<0.01 by Kruskal–Wallis test and Dunn’s multiple comparison test. (C) Locations for qPCR primers for the detection of 45S pre-rRNA, 18S rRNA and 28S rRNA. (D-F) TBL3 knock down increased 45S pre-rRNA/ACTB ratio but decreased 18S rRNA/45S pre-rRNA ratio, compared to siCtl treated cells. 45S pre-rRNA/ACTB, 18S rRNA/45S pre-rRNA and 28S rRNA/45S pre-rRNA ratios in siCtl treated cells were normalized to 1, respectively. Data were expressed as mean ± SEM from 3 independent samples per condition (N=3) for D-F. *p<0.05 and **p<0.01 by Kruskal–Wallis test and Dunn’s multiple comparison test. (G-I) 45S pre-rRNA/ACTB, 18S rRNA/45S pre-rRNA and 28S rRNA/45S pre-rRNA ratios in human control, HD and SCA2 postmortem cerebella. 45S pre-rRNA/ACTB, 18S rRNA/45S pre-rRNA and 28S rRNA/45S pre-rRNA ratios in Ctl group were normalized to 1, respectively. Data were expressed as mean ± SEM from N=3 (Ctl), 4 (HD) and 5 (SCA2) individual patient cerebella samples. *p<0.05 by Kruskal–Wallis test and Dunn’s multiple comparison test.

### 45S pre-rRNA level and processing is altered in SCA2 and HD postmortem tissue

Finally, we examined the expression of 45S pre-rRNA in human postmortem SCA2 and HD cerebella. qPCR amplification suggested that there was a slight, though not statistically significant, increase of 45S pre-rRNA level (normalized to *ACTB*) in both SCA2 and HD cerebella, compared with control (Fig. 4G). The qPCR results suggested that there was a decrease in both 18S rRNA maturation and 28S rRNA maturation in SCA2 and HD cerebella, compared with the controls (Fig. 4H-I), consistent with the trend observed with the knock down of TBL3 (Fig. 4D-F). Only the decrease in HD samples, but not in SCA2 samples, reached statistical significance, under the caveat that the relatively lower statistical power in SCA2 samples may not allow the detection of small changes. Taken together, the data supports the idea that aberrant RNA-RBP interactions may affect the steady state level and the maturation of 45S pre-rRNA in both SCA2 and HD.

## Discussion

We have previously shown that antisense *ATXN2-AS* transcripts contribute to SCA2 pathogenesis^42^. Here we provide the first evidence that sense *expATXN2* transcripts is involved in SCA2 pathogenesis. First, in establishing its potential pathogenicity, we show that untranslatable FL *ATXN2* transcript is neurotoxic (Fig. 1). This model is not suitable to test the contribution of sense *ATXN2* transcript relative to the toxicity of ATXN2 protein, or antisense *ATXN2-AS* RNA ^42^, because neither ATXN2 protein nor *ATXN2-AS* RNA is present in this model. In the future, genome editing approaches can be used to establish SCA2 iPS cell models that specifically model protein-versus RNA-triggered mechanism of pathogenesis. SCA2 iPSCs can also be subjected to transcriptome, proteome, and RNA interactome analysis to identify additional pathways that are involved in RNA-mediated aspects of SCA2 pathogenesis. Differentiation into neuronal types of greater and lesser selective vulnerability in SCA2 (e.g., Purkinje cells, cortical excitatory neurons, etc.) could be used to determine cell type vulnerability to RNA and protein mediated neurotoxicity. Next, we show that the *expATXN2* transcripts aggregate into nuclear RNA foci in SCA2 cell and transgenic mouse models, as well as in human SCA2 postmortem brain tissue. However, out of five human SCA2 postmortem brains available for this study, *expATXN2* RNA foci were only detected in case H1 that had the 22/38 *ATXN2* CAG repeat lengths and the latest disease on-set (Table 1). Interestingly, we have recently characterized a transcript that is expressed in the direction antisense to *ATXN2* (*ATXN2-AS*) and contains an expanded CUG repeat ^42^. CUG RNA foci containing this transcript were detected in SCA2 cases K3 and M1 (Table 1; ^42^). Given that SCA2 is associated with a relatively short repeat expansion, detection of foci may require a more sensitive assay. There are a number of alternative explanations for the absence of foci in the other SCA2 brains: (1) CAG RNA foci are highly toxic, or appear in cells marked for early death, such that Purkinje cells with foci may not have been present by the time of death; (2) CAG RNA foci are protective and associated with late onset disease and perhaps slower disease progression; (3) detectable foci were lost consequent to the process of brain collection or storage; (4) RNA foci are a byproduct of neurotoxic processes and have a neutral role in neurotoxicity; 5) RNA foci are an epiphenomenon, present in some SCA2 brains because of an unknown genetic or environmental factor and with no relevance to disease. Recent work describing a transgenic BAC mouse model expressing expanded *C9orf72* (*expC9orf72*) and exhibiting widespread RNA foci, but lacking behavioral abnormalities and neurodegeneration, even at advanced ages, suggests that RNA foci are not sufficient to trigger toxicity in ALS ^64^. A transgenic mouse model expressing non-translatable FL *ATXN2*, which could be tracked in live cells in real time ^65^ might help determine the relevance of CAG RNA foci to disease pathogenesis.

Our data strongly suggests that the neurotoxicity of *expATXN2* transcript involves aberrant *expATXN2*-RBP interactions that perturb rRNA maturation. We initially focused on Transducin beta like protein 3 (TBL3), a component of the SSU processome required for rRNA processing. While our RNA pull-down assay indicates that TBL3 interacts with *expATXN2* RNA, this assay cannot be used to prove a direct interaction. Filter-binding assays showed that recombinant TBL3 NTD can weakly interact with *expATXN2* RNA, and preferentially binds the structures of the CAG repeats. (Fig. 3D). Other components of the SSU processome likely stabilize the interaction of TBL3 with the *expATXN2* RNA in the cell. The yeast homolog of TBL3, Utp13, recognizes double-stranded regions of the pre-rRNA as a heterotetramer with other Utp proteins. Indeed, mass spectrometry analysis of *expATXN2* interactors did identify other proteins from the SSU processome in our isolated complexes (Table S1). One interesting possibility is that the multi-dentate recognition of structured RNA by TBL3 and its binding partners, which is a normal feature of their function in pre-rRNA processing, also contributes to the toxic aggregation of CAG repeat RNAs. Future experiments will be needed to determine which proteins are most important for neuronal toxicity.

Our results indicate that a subset of RBPs bind to both expanded *ATXN2* and *HTT* transcripts. This is not surprising, as it is well established that transcripts containing expanded *CAG* repeats form similar secondary structures *in vitro* ^35, 66, 67^ and, hence, at least some of the downstream effects are likely to be shared between different CAG repeat diseases. While SCA2 primarily affects cerebellum, HD is primarily characterized by atrophy of striatum and cerebral cortex ^68^. However, recent evidence indicates that cerebellum is also affected in HD ^69, 70^ and, in fact, appears to degenerate early ^71^ and independently from the striatal atrophy ^71^. This suggests that similar mechanism of pathogenesis may contribute to cerebellar pathology in both SCA2 and HD. Whether and to which degree mutant RNA-triggered mechanisms contribute to this pathology, remain to be further determined.

Interestingly, it was previously reported that expanded *ataxin-3* (*ATXN3*) transcripts, involved in spinocerebellar ataxia type 3 (SCA3) interact with nucleolin. In SCA3, this aberrant nucleolin-*ATXN3* interaction decreases 45S pre-rRNA levels in cell and *Drosophila* models of SCA3 ^31^. On the other hand, aberrant interaction between the *expC9orf72* transcripts and nucleolin, may contribute to the decreased maturation of 28S, 18S and 5.8S rRNAs from the precursor 45S pre-rRNA in ALS patients associated with CCCCGG hexamer expansion in *C9orf72* gene ^72^. It is quite possible that a therapeutic agent that prevents aberrant RNA-RBP interactions between toxic hairpin-forming transcripts and RBPs may be at least partially effective across multiple diseases. Alternatively, similar therapies may target shared pathogenic pathways downstream of the toxic transcripts.

In summary, we provide the first evidence that the *ATXN2* transcript with an expanded repeat may contribute to SCA2 pathogenesis, with similar properties to transcript-mediated toxicity in HD. The *ATXN2* transcript with an expanded CAG repeat itself, or its protein interactors, may provide valuable therapeutic targets in the future.

## Acknowledgments

We would like to acknowledge support for the statistical analysis from the National Center for Research Resources and the National Center for Advancing Translational Sciences (NCATS) of the National Institutes of Health through grant 1UL1TR001079. We thank Dr. Laura Ranum for the kind gift of the A8(*KKQ_EXP_)-3Tf1 construct. We thank Dr. Arnulf Koeppen, for providing frozen brain samples of three SCA2 patients. We thank Dr. Olga Pletinikova for providing a frozen brain sample of one SCA2 patient. We thank Dr. Shanshan Zhu for technical assistance regarding confocal imaging. We thank Kathryn A. Carson, Sc.M. for the advice on statistical analysis.

## Author contributions

D.D.R. and P.P.L conceived the study, oversaw the project, and designed the experiments; P.P.L., R.M., H.F., X.S., N.A., J.J., L.M., E.H., and D.D.R. carried out the experiments and analyzed data; H.Y.E.C., C.A.R., S.M.P., R.L.M., and S.W. provided fundamental reagents and intellectual contribution; P.P.L., R.M., R.L.M., S.W., and D.D.R. wrote the manuscript. D.D.R. contributed to this work prior to her current position. All the authors had final approval of the submitted version.

---

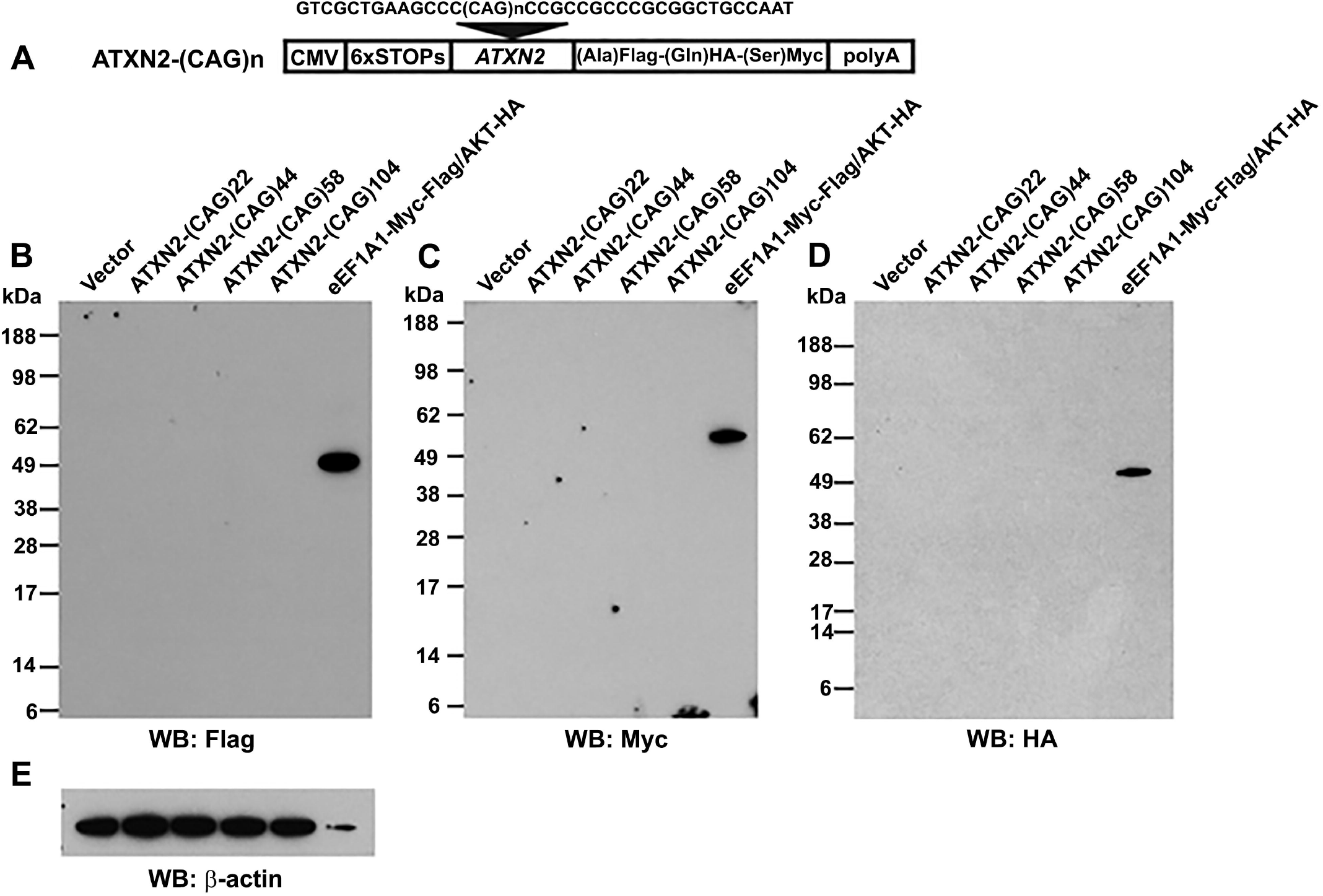

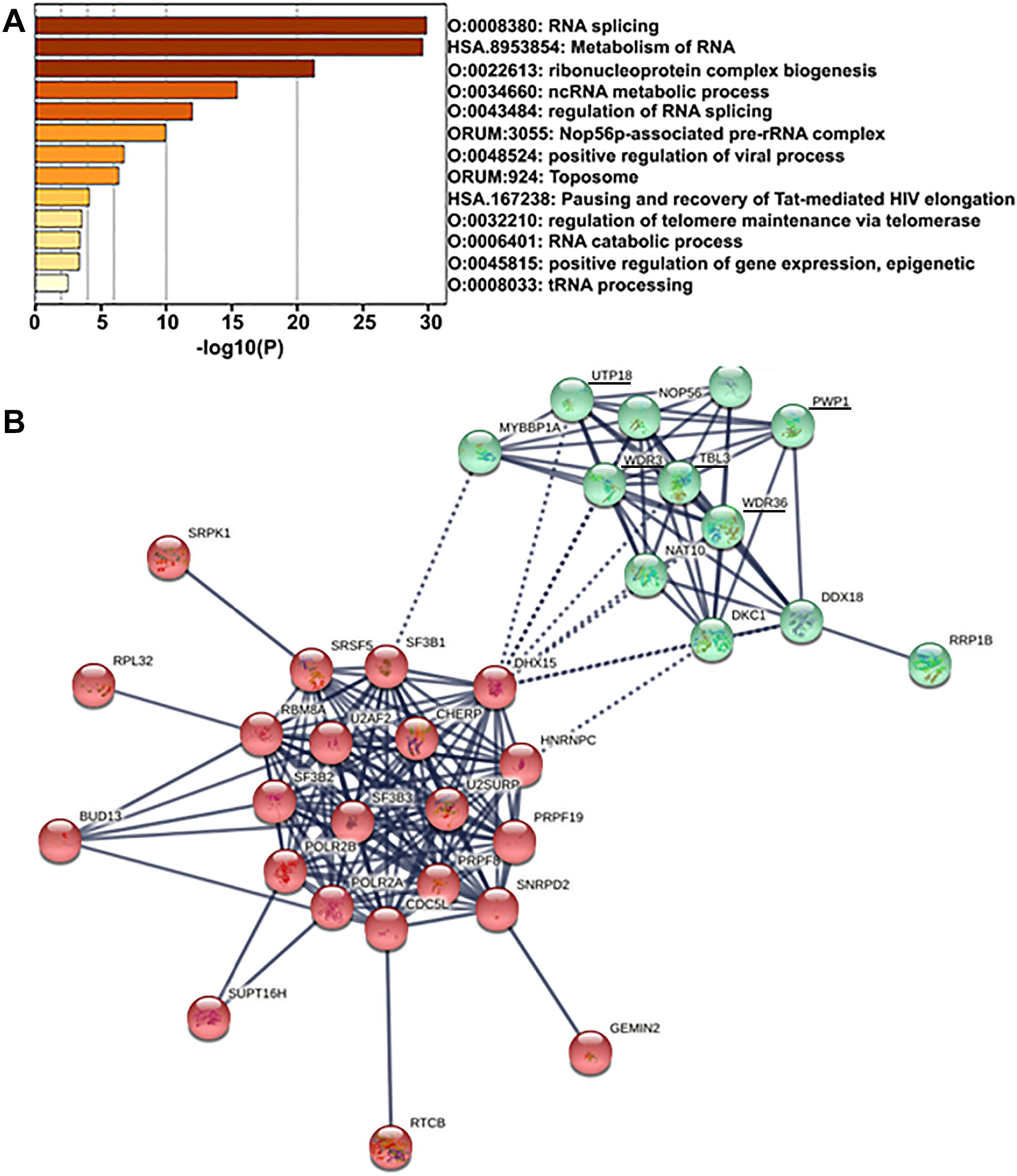

