## Supplementary data for "*RNA toxicity and perturbation of* rRNA processing in spinocerebellar ataxia type 2"

**Materials and Methods**

*Brain Tissue*

Five human SCA2, four HD, and three control brains were obtained from the Johns Hopkins University Brain Bank (Baltimore, MD), the University of Maryland Brain Bank (Baltimore, MD) and the hereditary ataxia tissue repository at the VA Medical Center (Albany, NY). Postmortem interval varied from 6 to 24 hours, and expanded allele lengths for SCA2 cases ranged from 37 to 44 triplets, while expanded allele lengths for HD cases ranged from 43 to 46. Control brains were matched to the SCA2 brains for age and postmortem interval. Clinical information for control, SCA2 and HD patients is included in Table 1. Wild-type (WT) and SCA2 transgenic (SCA2 Tg; ATXN2Q127) mice were maintained as previously described ^1^ following University of Utah IACUC and National Institutes of Health guidelines. Three-month-old female WT and SCA2 Tg mice were deeply anesthetized and cerebella collected. Tissues were kept at −80°C until the time of processing.

*Cell culture and transfection*

SK-N-MC neuroblastoma cells ^2^ and HEK293T cells (ATCC, Manassas, VA) were maintained in Dulbecco’s modified eagle medium with high glucose, supplemented with 10% fetal bovine serum and 1% penicillin/streptomycin/amphotericin B (Sigma). Plasmids were transfected into SK-N-MC cells using lipofectamine 2000 (Thermo Fisher) according to the manufacturer’s protocol. Primary mouse cortical neurons were prepared and transfected as previously described ^3^. Silencer select negative control #1 siRNA (undisclosed) and three TBL3 siRNAs (5’-GGACCAUCAAGAACAACGAtt-3’, 5’-CUCCGGCCGUGACAAGAUAtt-3’, 5’-GCCUUGCUGUCCAAGAACAtt-3’, Thermo Fisher) were reverse transfected into HEK293T cells using lipofectamine RNAiMAX reagent, as previously described ^4^.

*Plasmid constructs*

Full length (FL) N-terminal GFP-tagged ATXN2 (*GFP-ATXN2-Qn*) constructs were made by deleting *ATXN2* 1-480bp from the previously described GFP-ataxin-2 [Qn] constructs ^5^, resulting in translation initiation of the exogenous ATXN2 from the second ATG start codon, mimicking endogenous ATXN2 translation ^6^. Briefly, the GFP-ataxin-2 [Qn] was digested with Bgl2 and Xho1 to remove the ATXN2 1-1367bp, obtaining a GFP-ATXN2-Bgl2/Xho1 fragment. ATXN2 481-1367bp was PCR amplified from GFP-ataxin-2 [Qn] using primers 5’-GGCCGGACTCAGATCTCCATGTCGCTGAAGCCCCAG-3’ (forward) and 5’-TCCAGGGCCACTCGAGCTTTGTACTGGGCACTTGAC-3’ (reverse). The *GFP-ATXN2-Bgl2/Xho1* fragment and *ATXN2* 481-1367 fragment was ligated by infusion cloning (Takara) to obtain the final *GFP-ATXN2-Qn* constructs. The untagged *FL-ATXN2-Qn* constructs were made by subcloning the *FL-ATXN2-Qn* inserts from *GFP-ATXN2-Qn* into pcDNA3.1 vector. In order to obtain the FL non-translatable *Rluc-ATXN2-(CAG)n* constructs, a C-terminal Flag tagged Renilla luciferase (Rluc) cDNA sequence was cloned in between the Hind3 and Kpn1 sites (5’ of the *ATXN2* ORF) in *FL-ATXN2-Qn* constructs by infusion cloning and primers 5’-CTGGCTAGTTAAGCTATGACTTCGAAAGTTTATGA-3’ (forward) and 5’-GCTCGGTACCAAGCTTTTACTTATCGTCGTCATCCTTGTAATCTTGTTCATTTTTGAGAACTC-3’ (reverse). To obtain truncated ATXN2-Qn constructs, an *ATXN2* fragment containing the region 21 bp upstream of the CAG repeat and 105 bp downstream from the repeat was PCR-amplified using primers 5’-GCCACTGTGCTGGATATCCTCACCATGTCGCTGAAGCC-3’ (forward), 5’-TGGAATTCTGCAGATATCCGAGGACGAGGAGACCGAGG-3’ (reverse), and cloned into pcDNA3.1(-) myc-his A vector (Life Technologies) at EcoRV site. Normal control brain cDNA, and GFP-ataxin2 [Q104] plasmid ^5^ served as templates for the PCR. To examine RAN translation, *ATXN2-Qn* inserts were PCR-amplified and cloned into the XhoI and XbaI sites of A8(*KKQEXP)-3Tf1 vector, a kind gift from Dr. Laura P. W. Ranum to produce *ATXN2-(CAG)n* plasmids (Fig. 3). eEF1A1-Myc-Flag encoding myc- and flag-tagged eEF1A1 were from Origene, and HA-Akt DN (K179M) was a gift from Dr. Mien-Chie Hung (Addgene plasmid # 16243) ^7^. The *HTT-N63Q80* and *JPH3-(CTG)55* constructs were previously described ^8, 9^. The *ATXN2-AS-(CTG)104* construct was made by inverting the truncated *ATXN2-Q104* insert. The *HTT-AS-(CTG)80* construct was made by cloning *HTT-AS* exon1 with 80 CTG triplets into pcDNA3.1 vector at Hind3 and Xho1 sites ^10^.

*In vitro transcription and biotinylated RNA pull down*

As a first step in *in vitro* synthesis of *ATXN2-(CAG)n*, *ATXN2-AS-(CUG)104* and *JPH3-(CUG)55* RNA, truncated *ATXN2-Qn*, *ATXN2-AS-(CTG)104* and *JPH3-(CTG)55* plasmids were linearized with EcoR1. To synthesize *HTT-(CAG)n* and *HTT-AS-(CUG)80* transcripts *in vitro*, truncated *HTT N63Qn* and *HTT-AS-(CTG)n* constructs were linearized with Xho1. Biotinylated RNAs were *in vitro* transcribed from the linearized plasmids using MEGAscript (Thermo Fisher). In order to preserve the proper secondary structure of CAG or CUG repeats, biotin-16-UTP (Roche) was incorporated into *ATXN2-(CAG)n* and *HTT-(CAG)n* transcripts, while biotin-11-ATP (Perkin Elmer) was incorporated into *ATXN2-AS-(CUG)104*, *HTT-AS-(CUG)80* and *JPH3-(CUG)55* transcripts. 5 μg of biotinylated RNAs in structure buffer (10 mM Tris-HCl pH7.0, 0.1 M KCl and 10 mM MgCl_2_) were incubated at 70°C for 5 min, cooled on ice for 5 min, and then loaded with 40 μl of Dyna streptavidin beads (Life Technologies) according to the manufacturer’s protocol. To use Morpholinos (MO, Gene Tools) to block the CAG repeat region on the transcripts, 5 μg of biotinylated RNAs were mixed with 25 μg of either control (Ctl, 5’-CCTCTTACCTCAGTTACAATTTATA-3’) or (CTG)_8_C (5’-CTGCTGCTGCTGCTGCTGCTGCTGC-3’) MO ^11^ in structure buffer, incubated at 70°C and combined with the Dyna streptavidin beads. Neuronal-like SK-N-MC cells were lysed in buffer containing 25 mM Hepes pH7.4, 150 mM NaCl, 5mM MgCl2, 0.5% Triton, 40U/ml RNasin (Promega) supplemented with protease inhibitors (Sigma). SK-N-MC cell extract was incubated with Dyna beads loaded with different biotinylated RNAs for 1hr at 4°C. To wash off unbound proteins, beads were rinsed 5 times in cell lysis buffer for 10 min at 4°C. Proteins were eluted from Dyna beads using 0.1% SDS in H_2_O and submitted for MS analysis or used for western blotting. Rabbit anti-TBL3 antibody (HPA042562) was purchased from Sigma.

*Recombinant protein expression and purification*

To construct pMBP-TBL3 for bacterial expression of MBP tagged full-length human TBL3 protein, the full length TBL3 cDNA was cloned into the pMAL-c2G vector at the EcoRI and Hind III sites using NEBuilder HiFi DNA assembly cloning kit (NEB) and 5'-ACACTACGTAGAATTCATGGCAGAGACCGCGGCCGG-3' and 5'-GGCCAGTGCCAAGCTTCTAGGGCAGTGCGCCTTTAT-3' as the forward and reverse primers, respectively. To make pMBP-NTD-TBL3 expressing the TBL3 N-terminal domain (NTD, 1-633 aa), a stop codon was inserted after residue 633 using 5'- TAGTAGTAGAAGGATGTGACCGAGGCG-3' and 5'-CTACTACTACCAGAGGATGACTCGGGAGTCACT-3' as the forward and reverse primers, respectively.

BL21(DE3) cells transformed with pMBP-NTD-TBL3 were used to inoculate a 4-mL starter culture in Luria Broth (with 100 μg/mL ampicillin), which was grown for 10 h at 37 ˚C. The starter culture (2.5 mL) was used to inoculate 250 mL terrific broth (with 100 μg/mL ampicillin), grown with shaking at 37 ˚C to OD_600_ = 1, and then TLB3 expression was induced with 100 µg/mL isopropyl β-d-1-thiogalactopyranoside (IPTG) for 5 h. The cells were collected at 6,500 × *g* at 4 °C, and the cell pellet resuspended in 15 mL ice-cold Buffer A (50 mM Tris-HCl, 100 mM NaCl, 0.5% Triton X-100, 10 mM EDTA, 1 mL protease inhibitor (Roche), pH 8), sonicated (60 W with pulses of 5 s ON, 2 s OFF for 10 min) and spun at 31,000 × *g*. The pellet was resuspended twice in 15 mL Buffer B (50 mM Tris-HCl, 2.5 M guanidinium hydrochloride, pH 8), sonicated and spun as above. Finally, the cell pellet was resuspended in 40 mL Buffer C (50 mM Tris-HCl, 100 mM NaCl, 1 mM DTT, pH 8) containing 10 mM imidazole and 8 M urea (loading buffer), sonicated and spun as above. Hereafter, all steps were carried out at 4 °C. The supernatant was loaded on a 5-mL nickel-nitrilotriacetic acid agarose column pre-equilibrated with the loading buffer. The column was washed with 8 column volumes of loading buffer, and the protein was eluted with 30 mL Buffer C + 8 M urea + 40 mM imidazole, pH 8. The eluted fraction was refolded for 8 h by serial dropwise dilution; first 10-fold dilution in Buffer C + 0.2 M urea and then a two-fold dilution in Buffer C + 10 mM EDTA. The refolded protein was bound to amylose beads pre-equilibrated with Buffer C + 10 mM EDTA, washed with at least three column volumes of the same buffer, and the protein was eluted with Buffer C + 10 mM EDTA+ 20 mM maltose. The fractions containing eluted protein were pooled and dialyzed against 50 mM Tris-HCl, 10 mM EDTA, NaCl, 1 mM DTT, pH 8). The final solution was concentrated 10 to 15-fold using 15-mL concentrators (Millipore) and the purity was checked by SDS PAGE. The final concentration were determined using ε_280_=169760 M^-1^ cm^-1^ for MBP-NTD-TBL3 ^12^. The protein solution was flash frozen and stored at –80 ˚C.

To express full length recombinant muscle blind splicing regulator 1 (MBNL1, residues 1-382), the MBNL1 gene was synthesized and cloned into the BamHI and EcoRI sites in pGEX6P1, over-expressed in BL21DE3 cells and purified as previously described ^13^.

*Filter-binding assays*

Truncated *ATXN2* containing 22 or108 CAG repeats, or 105 interrupted CAA/CAG repeats, and *ATXN2-AS* containing 110 CUG repeats (see above) were transcribed in 250 µL total volume, using 5 µg EcoRI-linearized plasmid DNA template, 2 mM ATP, 2 mM UTP, 6 mM GTP, 6 mM CTP, 30 mM guanosine, 40 mM Tris-HCl (pH 8), 15 mM MgCl2, 5 mM DTT, 2 mM spermidine, T7 RNA polymerase at 37 °C for 4 h. The transcripts were purified by denaturing 6% PAGE and 5′-[^32^P] -labeled with T4 polynucleotide kinase and γ-[^32^P]-ATP (6000 Ci/mmol). The excess ATP was removed by passing the reaction through a TE-100 size exclusion column (Takara Biosciences). Prior to use in filter binding assays, the labelled RNAs were diluted to 1 nM (final) in 20 mM HEPES-K, 45 mM KCl, 15 mM NaCl, pH 7.4 and subjected to heat denaturation at 65 °C for 5 min followed by refolding at room temperature for 10 min. For binding assays, 100 pM of the 5’ end-labelled refolded RNA was incubated with 0-1500 nM MBP-NTD-TBL3 in 20 mM HEPES-K, 45 mM KCl, 15 mM NaCl, 1 mM DTT, 10% glycerol, 3 mM MgCl_2_ at pH 7.4 in the presence or absence of 0.1 mg/mL yeast tRNA, in a total volume of 20 µL. The RNA refolding and binding conditions were standardized using MBNL1 binding to the *expATXN2* transcripts as a positive control. Binding reactions were incubated at 4 °C for 2 h, then 18 µL of each reaction mixture was spotted on a microfiltration apparatus (BioRad) fitted with a double layer of nitrocellulose (Thermo Scientific) and Hybond-N plus (GE). Filters were washed once with binding buffer pre-chilled at 4 °C. The membranes were UV-crosslinked, air dried and imaged using a phosphor storage screen. After subtracting background from non-specific retention of RNA on the nitrocellulose membrane in the absence of protein, the amounts of ^32^P-labeled RNA on the nitrocellulose (NC) and Hybond membranes were quantified (ImageQuant) and the fraction bound, ƒ_B_, was calculated from ƒ_B_ = cpm NC/(cpm NC + cpm Hybond). The K_D_ was obtained from a fit versus ƒ_B_ to protein concentration to $f_{B}={Ax}/\left( x+K_{D} \right)$, in which x is protein concentration, and *A* is the maximum amount of RNA retained on the filter at saturating protein concentrations.

*RNA extraction and Real-time quantitative PCR (qPCR)*

RNA from transfected SK-N-MC cells and human postmortem cerebella were extracted by TRIZOL (Life Technologies), further purified (RNeasy Mini Kit, Qiagen) and cleaned of genomic DNA (Ambion Turbo DNA-free kit, Thermo Fisher). To measure the expression levels of 45S pre-rRNA, 18S rRNA and 28S rRNA in transfected SK-N-MC cells or human postmortem cerebella by qPCR, 500ng or 1μg of RNA was reverse transcribed using the ImProm-II Reverse Transcription System and random hexamer primers (Promega). PowerUp SYBR Green master mix (Thermo Fisher) was used for qPCR. 45S pre*-rRNA*, 18S rRNA, *28s* rRNA, and beta-actin (*ACTB*) primers were as previously described ^14, 15^. qPCR was performed using a QuantStudio 12K flex detection system (Thermo Fisher). Each qPCR experiment involving transfected cells was performed three (3) times. In each separate qPCR experiment, fresh RNA was extracted, converted to cDNA, and assayed in triplicate for 45S pre-rRNA, 18S rRNA, 28S rRNA, and *ACTB* expression. Each set of triplicates was averaged and normalized to the mean *ACTB* or 45S pre-rRNA of a control sample to yield normalized 45S pre-rRNA/*ACTB*, 18S rRNA/45S pre-rRNA or 28S rRNA/45S pre-rRNA expression. The means of normalized 45S pre-rRNA/*ACTB*, 18S rRNA/45S pre-rRNA and 28S rRNA/45S pre-rRNA for each construct from three (3) separate experiments were grouped (N=3) and subjected to final statistical analysis. The qPCR experiment involving human tissue was done in triplicate for each sample. Each set of triplicates was averaged and normalized to the mean *ACTB* or 45S pre-rRNA of a particular control brain to yield normalized 45S pre-rRNA/*ACTB*, 18S rRNA/45S pre-rRNA or 28S rRNA/45S pre-rRNA expression. The means of normalized 45S pre-rRNA/*ACTB*, 18S rRNA/45S pre-rRNA or 28S rRNA/45S pre-rRNA for each brain were grouped into control (N=3), HD (N=4), and SCA2 (N=5), and subjected to final statistical analysis, as describe below.

*Fluorescent in situ hybridization (FISH)*

The presence of foci containing *expATXN2* in human tissue, mouse tissue or transfected cells was detected using a 5’ Texas Red labeled 2-O-methyl CUG riboprobe (IDT, Coralville, IA) as previously described ^9^. FISH with DNase or RNase treatment was performed as previously described ^8^.

*Caspase 3/7 assay and nuclear condensation assay*

Cell viability was assessed by measuring caspase 3/7 activity (Caspase-Glo 3/7 Assay; Promega) in a 96-well format at 72 hours post-transfection, as previously described ^9, 15^. Each experiment was performed four (4) separate times by transfecting constructs into SK-N-MC cells with a different passage number. In each experiment, independent constructs were transfected in quadruplicates (four (4) technical replicates). For each quadruplicate, the mean value, normalized to control, was determined. For each construct, the mean values from four (4) separate experiments were averaged to provide a final outcome. Viability of transfected primary mouse cortical neurons was assessed by measuring the Hoechst staining intensity 48 hours post-transfection as previously described ^3^. The nuclear condensation assay was done by an experimenter blinded to the identity of constructs.

*Statistics*

Power calculation was based on preliminary results (not included in the final analysis) using 2-sample T test (PASS 13 ^16^; NCSS, LLC, Kaysville, UT). Samples of 3 and 4 for control and HD for human brain experiments are capable of detecting an effect size as low as 2.77 (statistical power = 0.80, alpha = 0.05), samples of 3 and 5 for control and SCA2 for the human brain experiments are capable of detecting an effect size as low as 2.67, and samples of 3 for each condition for the cell experiments are capable of detecting an effect size as low as 3.07 (statistical power = 0.80. alpha = 0.05). The small number of available brains (only 5 SCA2 postmortem brains were available to this study) was recognized as limiting the power to demonstrate the statistical significance of small differences in the data. The results were analyzed using nonparametric methods. Kruskal-Wallis test, followed by uncorrected Dunn multiple comparison test, was used for multiple-group comparison. Statistical significance was set at p < 0.05.

**Figure legends**

Fig. S1. Non-ATG initiated (RAN) translation does not contribute to the toxicity of *expATXN2* in SK-N-MC cells. (A) Schematic presentation of the *ATXN2-(CAG)n* constructs with the 6X STOP cassette and three tags (Flag, HA and myc) in three ORFs. (B-E) SK-N-MC cells were transfected with *ATXN2-(CAG)n* constructs, and the presence of RAN translation products in polyQ, polyalanine (polyAla) or polyleucine (polyLeu) ORFs was assessed by western blotting 72 hours post-transfection. pcDNA3.1 empty vector was used as a negative control. eEF1A1-Myc-Flag and Akt-HA plasmids were used as positive controls for antibodies used in the experiment. β-actin was used as a loading control. Note that positive controls were purposefully under-loaded. N=3 independent experiments and representative blots were shown.

Fig. S2. *expATXN2* RBPs grouped by functions. A total of 57 preferential *expATXN2* RBPs that were not identified in the bead only control, and were either identified as binding only to exp*ATXN2 (*with 58 and/or 104 repeats*)* or had at least twice the number of peptide hits in *ATXN2-(CAG)104* pull down compared to that in *ATXN2-(CAG)22*, were included in the analysis. (A) Go analysis of functional annotation. Preferential *expATXN2* interactors were submitted to Metascape (https://metascape.org/gp/index.html#/main/step1) for GO analysis of functional annotation. P value cut off was 0.05. -Log10(P) was shown. (B) , STRING network analysis using the STRING web server shows major subnetworks and potential protein–protein interactions among the 57 preferential *expATXN2* RBPs. Node colors represent different subnetworks based on k means clustering of 2. RBPs that are not connected within the network are not shown. The interaction score was set at 0.9 with the highest confidence. Two major clusters shown as pre-mRNA splicing (red) and ribosome biogenesis (green). Note that five proteins (TBL3, UTP18, WDR3, WDR36 and PWP2; underlined) are components of the small subunit (SSU) processome for ribosomal RNA processing.

Table S1. A selective list of RBPs that were identified as preferentially or specifically binding to *expATXN2* *in vitro* biotinylated RNA pull-down assay followed by mass spectrometry (MS).

| *expATXN2* RBPs | Characteristics and functions | Accession  number | MW  (kDa) | Peptides Identified | % Sequence Coverage |
| --- | --- | --- | --- | --- | --- |
| MYBBP1A | myb-binding protein 1A; transcriptional regulation | Q9BQG0 | 149 | 15 | 15% |
| RRP1B | ribosomal RNA processing protein 1 homolog B | Q14684 | 84 | 13 | 25% |
| CDC5L* | cell division cycle 5-like protein; cell cycle control, pre-mRNA splicing | Q99459 | 92 | 9 | 14% |
| NOP56 | nucleolar protein 56; rRNA biogenesis | O00567 | 66 | 9 | 20% |
| DDX50 | ATP-dependent RNA helicase DDX50 | Q9BQ39 | 83 | 6 | 12% |
| ZC3H18 | zinc finger CCCH domain-containing protein 18 | Q86VM9 | 106 | 6 | 9% |
| KNOP1 | Lysine-rich nucleolar protein 1 | Q1ED39 | 52 | 5 | 14% |
| TBL3*^ | transducin beta-like protein 3 | Q12788 | 89 | 5 | 8% |
| NKRF | NF-kappa-B-repressing factor; transcriptional regulation | O15226 | 79 | 5 | 7% |
| WDR36*^ | WD repeat-containing protein 36 | Q8NI36 | 105 | 4 | 5% |
| RTCB* | tRNA-splicing ligase RtcB homolog | Q9Y3I0 | 55 | 4 | 8% |
| SPATS2L | SPATS2-like protein | Q9NUQ6 | 62 | 4 | 9% |
| ZC3H4* | zinc finger CCCH domain-containing protein 4 | Q9UPT8 | 140 | 4 | 7% |
| FXR2 | fragile X mental retardation syndrome-related protein 2 | P51116 | 74 | 4 | 11% |
| UTP18*^ | U3 small nucleolar RNA-associated protein 18 homolog | Q9Y5J1 | 62 | 4 | 8% |
| mH2A1* | core histone macro-H2A.1; chromatin modification | O75367 | 39 | 3 | 10% |
| DDX18 | ATP-dependent RNA helicase DDX18 | Q9NVP1 | 75 | 3 | 5% |
| NKAP* | NF-kappa-B-activating protein; transcriptional regulation | Q8N5F7 | 47 | 3 | 10% |
| CHD-4* | chromodomain-helicase-DNA-binding protein 4; histone deacetylation | Q14839 | 218 | 3 | 2% |
| CHERP | calcium homeostasis endoplasmic reticulum protein; calcium homeostasis | Q8IWX8 | 104 | 3 | 4% |
| pNO40* | Nucleolar protein of 40 kDa; | Q9NP64 | 28 | 2 | 12% |
| Aip-4 | ADP-ribosylation factor-like protein 6-interacting protein 4; pre-mRNA splicing | Q66PJ3 | 36 | 2 | 8% |
| U2AF65* | Splicing factor U2AF 65 kDa subunit; pre-mRNA splicing, mRNA export | P26368 | 54 | 2 | 4% |
| DHX57* | putative ATP-dependent RNA helicase DHX57 | Q6P158 | 156 | 2 | 2% |
| hPOP1 | ribonucleases P/MRP protein subunit POP1 | Q99575 | β-like protein 3 (TBL-3)115 | 2 | 2% |
| FAM133B* | protein FAM133B | Q5BKY9 | 28 | 2 | 13% |
| SCAF* | splicing factor, arginine/serine-rich 19; pre-mRNA splicing | Q9H7N4 | 139 | 2 | 2% |
| MRP-L47 | 39S ribosomal protein L47, mitochondrial | Q9HD33 | 29 | 2 | 8% |
| WDR3*^ | WD repeat-containing protein 3 | Q9UNX4 | 106 | 2 | 4% |
| PWP1^ | periodic tryptophan protein 1 homolog | Q13610 | 56 | 2 | 3% |
| BUD13* | BUD13 homolog | Q9BRD0 | 71 | 2 | 3% |
| RBBP6* | E3 ubiquitin-protein ligase RBBP6; protein ubiquitination | Q7Z6E9 | 197 | 2 | 2% |
| POLR2B* | DNA-directed RNA polymerase II subunit RPB2; transcriptional regulation | P30876 | 134 | 2 | 1% |

*RBPs binding to *expHTT* transcript identified by MS as well; ^components of SSU processome.
